# Does vendor breeding colony influence sign- and goal-tracking in Pavlovian conditioned approach? A preregistered empirical replication

**DOI:** 10.1101/2022.07.09.499421

**Authors:** Shaun Yon-Seng Khoo, Alexandra Uhrig, Anne-Noël Samaha, Nadia Chaudhri

## Abstract

Vendor differences are thought to affect Pavlovian conditioning in rats. After observing possible differences in sign-tracking and goal-tracking behaviour with rats from different breeding colonies, we performed an empirical replication of the effect. 40 male Long-Evans rats from Charles River colonies ‘K72’ and ‘R06’ received 11 Pavlovian conditioned approach training sessions (or “autoshaping”), with a lever as the conditioned stimulus (CS) and 10% sucrose as the unconditioned stimulus (US). Each 58-min session consisted of 12 CS-US trials. Paired rats (n = 15/colony) received the US following lever retraction. Unpaired control rats (n = 5/colony) received sucrose during the inter-trial interval. Next, we evaluated the conditioned reinforcing properties of the CS, by determining whether rats would learn to nose-poke into a new, active (vs. inactive) port to receive CS presentations alone (no sucrose). Preregistered confirmatory analyses showed that during autoshaping sessions, Paired rats made significantly more CS-triggered entries into the sucrose port (i.e., goal-tracking) and lever activations (sign-tracking) than Unpaired rats did, demonstrating acquisition of the CS-US association. Confirmatory analyses showed no effects of breeding colony on autoshaping. During conditioned reinforcement testing, analysis of data from Paired rats alone showed significantly more active vs. inactive nosepokes, suggesting that in these rats, the lever CS acquired incentive motivational properties. Analysing Paired rats alone also showed that K72 rats had higher Pavlovian Conditioned Approach scores than R06 rats did. Thus, breeding colony can affect outcome in Pavlovian conditioned approach studies, and animal breeding source should be considered as a covariate in such work.

## Introduction

In Pavlovian conditioned approach (or “autoshaping”) studies, an environmental cue that is repeatedly paired with a reinforcer can acquire incentive salience [1–12]. During this process, motivational value becomes assigned to the cue and this is thought to play an important role in psychological disorders such as substance use disorder [13–15]. Animals in Pavlovian conditioned approach studies are often classified as sign-trackers, based on their propensity to interact with the cue, or goal-trackers, based on their preference for approaching the location of reinforcer presentation. Animals can also acquire an intermediate phenotype when they do not display a clear preference for sign-tracking or goal-tracking [8].

Across several Pavlovian conditioned approach experiments in rats (including [12] and unpublished results), we observed what appeared to be vendor differences in sign- and goal-tracking behaviours. We calculated Pavlovian conditioned approach (PCA) scores, which are typically defined as indicating sign-tracking when ≥ 0.5 and goal-tracking when ≤ −0.5 [8]. In some experiments, animals tended towards sign-tracking with mean PCA scores of approximately 0.3, while other cohorts of animals more frequently acquired a goal-tracking phenotype with mean PCA scores of approximately −0.3. Rats had been purchased from the same supplier but originated from one of two breeding colonies. Examining data across experiments suggested that rats from one colony (K72, Kingston, NY, United States) tended to become sign-trackers more often than rats from the other (R06, Raleigh, NC, United States).

Vendor differences have been reported in the literature since at least 1968, when Miller and colleagues reported different basal serotonin and noradrenaline levels in Sprague-Dawley rats from different suppliers [16]. In 1984, Sparber and Fossom first described within-vendor behavioural differences [17]. Sparber and Fossom found that rats seemed to differ in their operant response rates for food following amphetamine administration and learned that, despite being sourced from the same supplier, their rats came from different breeding colonies operated by the same company. They found that when they directly compared rats from the two colonies, they differed in their rate of acquisition of operant responding for food [17], indicating differences in learning and/or motivation. Since these two early papers, more than two dozen other papers have been published demonstrating between and/or within-vendor differences in rats on physiological (Table 1) or behavioural measures (Table 2). Similarly, behavioural effects due to vendor and substrain have been reported in mice since at least as early as 1972 when Poley reported C57BL/6J mice, but not C57BL/6A mice, preferred 10% alcohol to water [18]. Bryant and colleagues tabulated over a dozen studies up to 2008 that demonstrated behavioural differences in mice across different substrains [19]. Bryant and colleagues also reported data showing differences in rotarod performance and pain sensitivity across substrain and vendor [19]. Since then, additional reports of behavioural differences across vendors have continued to emerge [20–22].

**Table 1.** Papers reporting vendor differences in neurophysiology in rats

| Animals | Study Type | Effects | Reference |
| --- | --- | --- | --- |
| Male Sprague–Dawley rats | Experimental | Between–vendor: Basal serotonin norepinephrine | Miller et al. (1968) [16] |
| Male Sprague–Dawley rats | Experimental | Between–vendor: Basal serotonin norepinephrine | Sparber & Fossum (1984) [17] |
| Male Wistar and Sprague–Dawley rats | Experimental | Between–vendor: Focal cerebral ischemia | Oliff et al. (1995) [31] |
| Male Wistar and Sprague–Dawley rats | Experimental | Between–vendor: Collateral Anastomoses | Oliff et al. (1997) [32] |
| Male Sprague–Dawley rats | Experimental | Between–vendor: Bodyweight, NO synthase inhibition effect | Pollock et al. (1998) [33] |
| Male Wistar rats | Experimental | Between–vendor: Melatonin secretion | Barassin et al. (1999) [34] |
| Pregnant female Sprague–Dawley rats | Experimental | Between–vendor: NO synthase inhibition effect | Buhimschi et al. (2001) [35] |
| Male Sprague–Dawley rats | Experimental | Between–vendor: Motor response to hypoxia | Fuller et al. (2001) [36] |
| Female Sprague–Dawley and Wistar rats | Experimental | Between–vendor: MK-801 neurotoxic effects | Bueno et al. (2003) [37] |
| Wistar rats | Experimental | Between–vendor: Acute ischemia | Marosi et al. (2006) [38] |
| Male Sprague–Dawley rats | Experimental | Between–vendor: Metabolism, metabolic response to stress | Pecoraro et al. (2006) [39] |
| Wistar–Kyoto rats and Spontaneously Hypertensive Rats | Experimental + consortium data | Between–vendor: Genetic architecture | Zhang–James et al. (2013) [40] |

**Table 2.**
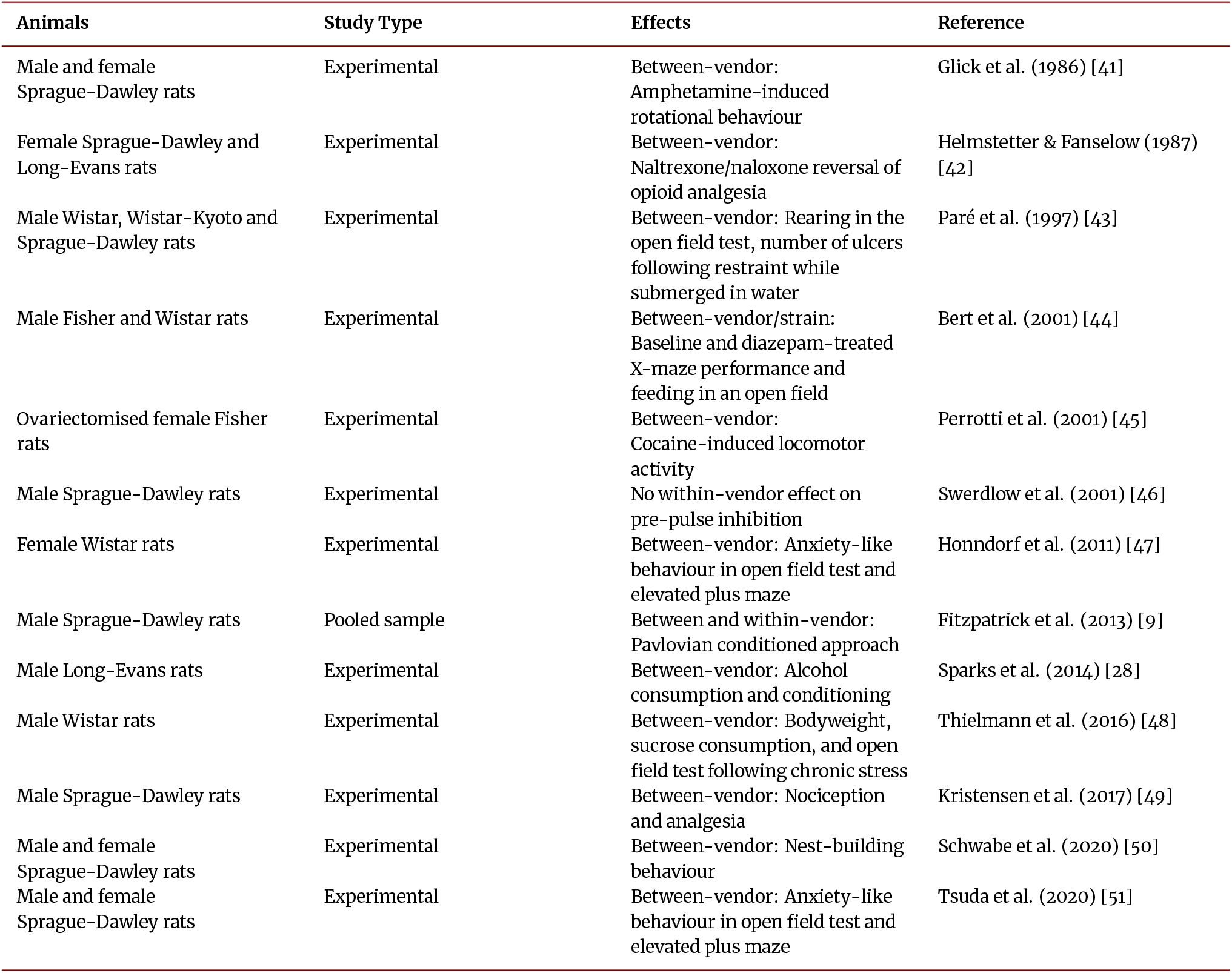
Papers reporting vendor differences in behaviour or behavioural pharmacology in rats

| Animals | Study Type | Effects | Reference |
| --- | --- | --- | --- |
| Male and female Sprague-Dawley rats | Experimental | Between-vendor: Amphetamine-induced rotational behaviour | Glick et al. (1986) [41] |
| Female Sprague-Dawley and Long-Evans rats | Experimental | Between-vendor: Naltrexone/naloxone reversal of opioid analgesia | Helmstetter & Fanselow (1987) [42] |
| Male Wistar, Wistar-Kyoto and Sprague-Dawley rats | Experimental | Between-vendor: Rearing in the open field test, number of ulcers following restraint while submerged in water | Paré et al. (1997) [43] |
| Male Fisher and Wistar rats | Experimental | Between-vendor/strain: Baseline and diazepam-treated X-maze performance and feeding in an open field | Bert et al. (2001) [44] |
| Ovariectomised female Fisher rats | Experimental | Between-vendor: Cocaine-induced locomotor activity | Perrotti et al. (2001) [45] |
| Male Sprague-Dawley rats | Experimental | No within-vendor effect on pre-pulse inhibition | Swerdlow et al. (2001) [46] |
| Female Wistar rats | Experimental | Between-vendor: Anxiety-like behaviour in open field test and elevated plus maze | Honndorf et al. (2011) [47] |
| Male Sprague-Dawley rats | Pooled sample | Between and within-vendor: Pavlovian conditioned approach | Fitzpatrick et al. (2013) [9] |
| Male Long-Evans rats | Experimental | Between-vendor: Alcohol consumption and conditioning | Sparks et al. (2014) [28] |
| Male Wistar rats | Experimental | Between-vendor: Bodyweight, sucrose consumption, and open field test following chronic stress | Thielmann et al. (2016) [48] |
| Male Sprague-Dawley rats | Experimental | Between-vendor: Nociception and analgesia | Kristensen et al. (2017) [49] |
| Male and female Sprague-Dawley rats | Experimental | Between-vendor: Nest-building behaviour | Schwabe et al. (2020) [50] |
| Male and female Sprague-Dawley rats | Experimental | Between-vendor: Anxiety-like behaviour in open field test and elevated plus maze | Tsuda et al. (2020) [51] |

If Pavlovian conditioned approach is similarly affected by between or within vendor effects, then this would have important implications. Pavlovian conditioned approach studies often examine shifts in the phenotypes of animals between sign- and goal-tracking [23–26]. These kinds of studies would therefore need to include consideration of vendor effects, especially if they are run using multiple cohorts or across multiple laboratories. More care would also be necessary when making comparisons across studies, since there may be baseline differences in the propensity to signor goal-track in rats from different vendors or colonies.

There is evidence in the Pavlovian conditioning literature that vendor differences influence phenotype. Kehoe and colleagues reported vendor differences in acquisition of a cue-conditioned reflex in rabbits [27]. Sparks and colleagues have also reported vendor differences in home-cage alcohol consumption and port entries triggered by an alcohol cue in Long-Evans rats [28]. Finally, Fitzpatrick and colleagues conducted a large analysis of rats from multiple projects and found both vender and breeding colony effects [9]. They pooled Pavlovian conditioned approach data from Sprague Dawley rats, with 115 rats from 2 Charles River colonies and 442 rats from 3 Harlan colonies (now Envigo). When they analysed the effect of breeding colony on sign-tracking and goal-tracking phenotypes after 5 days of autoshaping, they found significant effects both between and within vendors [9]. However, unlike the majority of studies on vendor differences (Tables 1 and 2), Fitzpatrick and colleagues did not directly compare rats from different vendors within a single experiment. It is therefore important to confirm whether within-vendor differences have relevance in Pavlovian conditioned approach studies by experimentally replicating their findings.

Based on the studies described above, we hypothesised that rats from different breeding colonies but supplied by the same vendor were differentially predisposed towards sign- and goal-tracking. Prior to conducting this study, we preregistered our hypotheses, design, and analysis plan [29]. Preregistration allowed us to transparently record our hypotheses, design, and analysis plan prior to data collection and analysis [30]. Our preregistered hypotheses were: (1) after autoshaping, K72 rats will have higher (more positive) Pavlovian Conditioned Approach scores (PCA scores; see Table 3) than rats from R06, (2) a greater proportion of K72 rats will meet criteria for classification as sign-trackers (PCA Score ≥ 0.5), and (3) rats from K72 will make more active nosepokes than rats from R06 in a conditioned reinforcement test. While we did not find support for any of these hypotheses in our preregistered confirmatory analyses, exploratory analyses that focussed on Paired rats alone provided some support for an effect of breeding colony.

**Table 3.** Calculation of the Pavlovian Conditioned Approach score and its components

| Score | Calculation |
| --- | --- |
| Response Bias | (Lever Activations - Port Entries) ÷ (Lever Activations + Port Entries) |
| Latency Score | (Mean Port Entry Latency - Mean Lever Activation Latency) ÷ CS Duration |
| Probability Difference | (Trials with Lever Activations - Trials with Port Entries) ÷ Number of Trials |
| PCA score | (Response Bias + Latency Score + Probability Difference) ÷ 3 |

## Methods

### Animals

A total of 40 male Long-Evans rats were obtained from Charles River (Kingston, NY & Raleigh, NC, United States). 20 rats were from the K72 (Kingston, NY) breeding colony and 20 rats were from R06 (Raleigh, NC). Sample size was based on a power analysis using G*Power 3.1.9.2 [52], which indicated that a total of at least 36 rats was required to achieve 95% power to detect an effect size of *ηp*^2^ = 0.09 with α of 0.05, 4 groups, 10 measurements, a correlation among repeated measures of 0.5, and non-sphericity correction of ε = 0.4. The effect size was based on unpublished observations between experiments using K72 and R06 rats. The majority of rats were allocated to the Paired condition (n = 15 per colony) because it is already well established that Unpaired rats do not acquire a conditioned response [24]. Rats were housed in standard wire-top plastic cages (44.5 cm 25.8 cm 21.7 cm) with Teklad Sani chip bedding (Cat# 7090, Envigo, Quebec, Canada) and unrestricted access to food (Teklad, Envigo, QC, Canada) and water. Rats were pair-housed on arrival and then individually housed after 3 days to enable measurement of home-cage sucrose consumption. Animals remained singly-housed to keep housing conditions consistent throughout the experiment. All procedures were approved by the Concordia University Animal Research Ethics Committee and conducted in accordance with guidelines from the Canadian Council on Animal Care.

### Apparatus

Behavioural training was conducted in 12 Med Associates (Fairfax, VT, United States) extra tall modular test chambers. Each chamber was contained in a sound attenuating cubicle with a fan to mask external noise. The fans were switched on whenever rats were brought into the behavioural testing room. A white houselight was situated in the centre of the left wall, near the ceiling of the chamber. On the right wall, there was a fluid port with infrared detectors located above the floor, flanked on either side by retractable levers. Levers were calibrated so that they could be activated by a 26-g weight. The fluid port was connected to a 3.3 RPM syringe pump which would be loaded with a 20 mL syringe for sucrose delivery.

### Home-cage sucrose

All rats were first familiarised with 10% sucrose (w/v). During this time they received 48 h of unrestricted access to a 90 mL sucrose bottle and a standard bottle of water. The sucrose and water bottles were weighed before access, then re-weighed and refilled after 24 h and then re-weighed after a second 24 h period to determine consumption. Spillage was accounted for using two empty control cages with sucrose and water bottles.

### Habituation

Rats were first habituated to transport from the colony room to the behavioural testing room. Rats were transported to the behavioural testing room, handled, weighed and then returned to their home-cage. After 20 min, they were transported back to the colony room.

The next day, rats were habituated to the conditioning chambers. Rats were placed in the conditioning chambers and, after a 2 min delay, the houselight was switched on for the duration of the 20 min session. During this session, port entries were recorded but had no programmed consequences.

### Pavlovian Autoshaping

Rats then received Pavlovian autoshaping. Autoshaping was planned to run for 10-14 sessions until mean PCA scores varied by less than 10% across two days, which occurred after 11 sessions. Each session lasted a total of 58 min and had 12 × 30 s trials, which consisted of a 10 s Pre-CS period, a 10 s CS presentation, followed by a 10 s Post-CS period. Trial onset was synchronised across boxes. For rats in the Paired group (K72 n = 15, R06 n = 15), the syringe pump was run for the first 6 s of the Post-CS period, to deliver 0.2 mL of 10% sucrose. Chambers were counterbalanced to present either the left or the right lever as the cue. The inter-trial interval (ITI) was randomly delivered as 240 ± 120 s. For Unpaired rats (K72 n = 5, R06 n = 5), sucrose delivery occurred during the ITI to equate total sucrose exposure in both Paired and Unpaired rats. Port entries during the CS are presented as a difference score (ΔCS port entries = CS port entries – Pre-CS port entries) to remove baseline levels of responding [12, 53].

For each session, the PCA score and its components were calculated according to the formulae described by Meyer and colleagues [8]. The PCA score is calculated as the mean of its three component measures, response bias, latency score, and probability difference (Table 3). Phenotypic classification was based on the mean PCA score from the final two sessions [8, 54]. Rats with a score ≥ 0.5 were classified as sign-trackers, ≤ −0.5 as goal-trackers, and the remaining rats were classified as intermediates.

### Conditioned Reinforcement

After Pavlovian autoshaping, the chambers were reconfigured to remove the levers on either side of the port and replace them with nosepokes. The port was then replaced with a retractable lever. First, rats were habituated to the nosepokes in a 10 min session. During this habituation sessionresponses were recorded but had no programmed consequences.

The following day, rats were tested in a 60 min conditioned reinforcement session. One nosepoke (counterbalanced) was designated the active nosepoke and would deliver a 2.5-s presentation of the lever when response ratios were met. The other nosepoke was designated as inactive and had no programmed consequences. The first 3 lever presentations were available on an FR1 schedule. Subsequent lever presentations were available on a VR2 schedule, requiring 1, 2, or 3 active nosepokes per presentation.

### Statistical Analysis and Material Availability

Data was analysed using SPSS 26 (IBM, Armonk, NY, United States). Preregistered confirmatory analyses were mixed-design ANOVAs with autoshaping Session as within-subjects factors and between-subjects factors of Colony (K72 vs R06) and Pairing (Paired vs Unpaired). A χ^2^ test was planned to test whether the proportion of sign-trackers and goal-trackers differed between Paired K72 and Paired R06 rats. For the conditioned reinforcement test, preregistered confirmatory analyses were a mixed-design ANOVA with Nosepoke (active vs inactive) as the within-subjects factor and between-subjects factors of Colony and Pairing and, for the number of lever presentations and activations, a 2×2 ANOVA with Colony and Pairing as between-subjects factors. When Mauchly’s test of Sphericity was significant, a Greenhouse-Geisser correction was applied to degrees of freedom for ε < 0.75. Post-hoc tests were Bonferroni-corrected.

Exploratory analyses were performed to analyse Paired rats alone, as done by Fitzpatrick and colleagues [9]. Linear mixed modelling of PCA Score was performed with an autoregressive (AR1) covariance structure [9] and maximum likelihood estimation. Beginning with the null model, the fixed factors of Colony and Session were added to the model, with model selection based on Hurvich and Tsai’s information criterion (AICc). Hurvich and Tsai’s information criterion was chosen because it corrects Akaike’s Information Criteria (AIC) for use with small samples [55], while other commonly used information criteria (e.g. AIC and Schwarz’s Bayesian Criterion) are most appropriate when sample sizes are greater than 100 [56]. Models were ranked and selected based on change in AICc (Δi), Akaike weight (w_*i*_), and evidence ratio (ER) [57]. To compare the results of the present study with those of Fitzpatrick and colleagues [9], 90% confidence inter-vals of effect size (η*p*^2^) were calculated from the fixed effects test using the apaTables package in R 4.1.2 [58].

Raw data underlying figures and Med-PC code is available on Zenodo [59].

## Results

### Paired rats acquired the CS-US association

Over the course of 11 autoshaping sessions there were overall increases in the number of CS lever activations (Figure 1a; F(3.372,121.406) = 3.006, p = 0.028, ε = 0.337). Although inspection of Figure 1a suggests that compared to Unpaired rats, Paired rats appear to activate the lever CS more frequently across sessions, there were no significant effects of Pairing (F(1,36) = 2.058, p = 0.16) and the Session × Pairing interaction was not quite significant (F(3.372,121.406) = 2.519, p = 0.054, ε = 0.337). There was no significant Pairing × Colony interaction (F(1,36) = 0.954, p = 0.335). There was also no significant main effect of Colony (F(1,36) = 0.395, p = 0.534) and no other interactions were significant (Session × Colony: F(3.372,121.406) = 0.879, p = 0.464, ε = 0.337; Session × Pairing × Colony: F(3.372,121.406) = 0.277, p = 0.864, ε = 0.337). Thus, rats generally made an increasing number of contacts with the lever CS over the course of autoshaping and there were no statistically significant differences between Paired and Unpaired rats in this effect.

**Figure 1.**
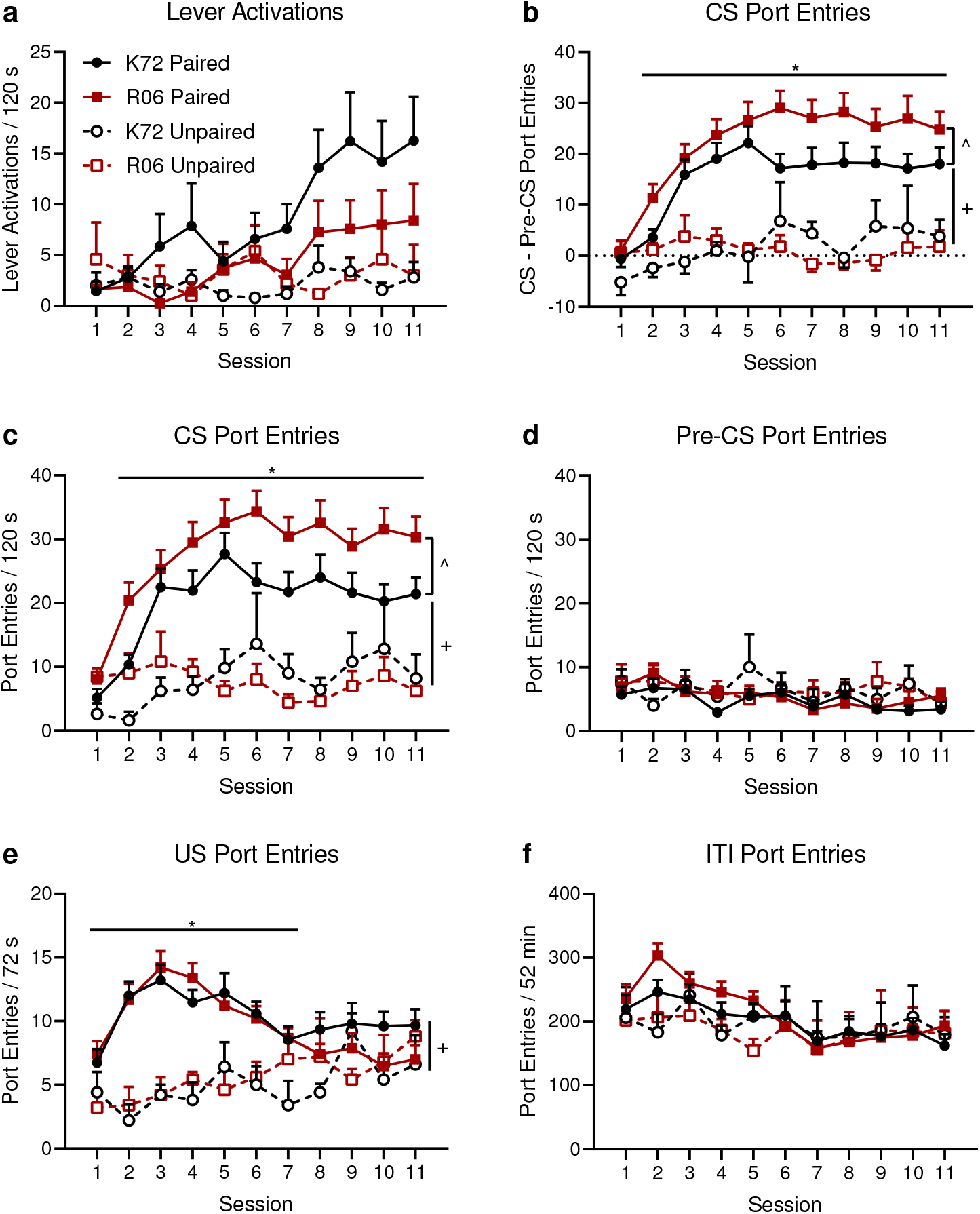
(a) Rats made more lever activations over the course of autoshaping. (b) Paired rats made significantly more ΔCS port entries (CS port entries – Pre-CS port entries) than Unpaired rats did in sessions 2-11. (c) Paired rats made significantly more unadjusted CS port entries relative to Unpaired rats in sessions 2-11. (d) Paired and Unpaired rats made similar numbers of pre-CS port entries across training. (e) Paired rats made significantly more US port entries than Unpaired rats did earlier in training (sessions 1-7). The US occurred immediately after CS presentation for Paired rats and during the ITI for Unpaired rats. (f) ITI port entries did not differ between Paired and Unpaired rats. * p < 0.05 in Bonferroni-corrected post-hoc comparisons. + p < 0.05 for main effect of Pairing. K72 Paired n = 15, R06 Paired n = 15, K72 Unpaired n = 5, R06 Unpaired n = 5. ^ When Paired rats were analysed alone, main effects of colony were observed for ΔCS and unadjusted CS port entries.

Analysis of both CS-triggered port entries (Figure 1b) and unadjusted CS port entries (Figure 1c) showed that pairing the CS and US influenced behaviour during autoshaping. Only Paired rats significantly increased their CS-triggered port entries over the course of autoshaping. ΔCS port entries, which subtract baseline Pre-CS port entries from CS port entries, significantly increased with Session (F(4.556,164.008) = 8.309, p < 0.001, ε = 0.456; Figure 1b). Overall ΔCS port entries were higher in Paired versus Unpaired rats, as shown by a significant main effect of Pairing (F(1,36) = 35.165, p < 0.001). Moreover, Paired rats increased ΔCS port entries with more training relative to Unpaired rats, as indicated by a significant Session × Pairing interaction (F(4.556,164.008) = 4.19, p = 0.002, ε = 0.456). Bonferonni-corrected post-hoc comparisons for this interaction indicated that Paired rats made more ΔCS port entries than Unpaired rats did in sessions 2-11 (p ≤ 0.008). However, there was no significant effect of Colony (F(1,36) = 1.179, p = 0.285) or Pairing × Colony interaction (F(1,36) = 1.668, p = 0.205). There was also no significant Session × Colony interaction (F(4.556,164.008) = 0.205, p = 0.951, ε = 0.456) or Session × Pairing × Colony interaction (F(4.556,164.008) = 1.271, p = 0.281, ε = 0.456). Thus, only Paired rats acquired a CS-triggered Pavlovian conditioned approach response to the sucrose port, and there were no significant effects of breeding colony.

Unadjusted CS port entries (Figure 1c) showed the same pattern as ΔCS port entries. CS port entries significantly increased with Session (F(4.74,170.657) = 7.17, p < 0.001, ε = 0.474) and were also significantly higher in Paired rats compared to Unpaired rats (Pairing: F(1,36) = 36.512, p < 0.001). Paired rats increased CS port entry responding relative to Unpaired rats, as shown by a significant Session × Pairing interaction (F(4.74,170.657) = 4.057, p = 0.002, ε = 0.474). As with ΔCS port entries, Bonferroni-corrected post-hoc comparisons indicated that, apart from session 1 (p = 0.533), Paired rats made more CS port entries in sessions 2-11 (p ≤ 0.002). However, there was no significant effect of breeding Colony (F(1,36) = 1.82, p = 0.186) or Pairing × Colony interaction (F(1,36) = 2.31, p = 0.137). There was also no Session × Colony interaction (F(4.74,170.657) = 0.492, p = 0.772, ε = 0.474) or Session × Pairing × Colony interaction (F(4.74,170.657) = 1.198, p = 0.313, ε = 0.474). The consistency between unadjusted and ΔCS port entries further highlights that Paired rats reliably acquired CS-triggered Pavlovian conditioned approach to the sucrose port.

Pre-CS port entries reflect baseline levels of entry into the sucrose port and as expected, responding was unchanged over the course of autoshaping (Figure 1d). Mixed-design ANOVA found no main effect of Session (F(6.951,250.23) = 1.654, p = 0.122, ε = 0.695). There was no overall difference between Paired and Unpaired rats (Pairing: F(1,36) = 2.056, p = 0.16) and no Session × Pairing interaction (F(6.951,250.23) = 0.873, p = 0.528, ε = 0.695). There was also no main effect of Colony (F(1,36) = 0.242, p = 0.626), Pairing× Colony interaction (F(1,36) = 0.104, p = 0.749), Session × Colony interaction (F(6.951,250.23) = 0.939, p = 0.476, ε = 0.695), or Session × Pairing × Colony interaction (F(6.951,250.23) = 0.577, p = 0.773, ε = 0.695). Thus, baseline levels of entry into the sucrose port did not change across autoshaping.

Port entries made during the 6 s of sucrose delivery (US port entries) also differed between Paired and Unpaired rats (Figure 1e). For Paired rats, US delivery began immediately after the CS, while for Unpaired rats US delivery was not signalled and occurred during the ITI. Mixed-design ANOVA indicated that there were overall increases in US port entries across Sessions (F(7.086,255.088) = 2.447, p = 0.019, ε = 0.709) and that overall US port entries was greater in Paired rats relative to Unpaired rats (Pairing: F(1,36) = 25.106,p < 0.001). The trajectory of the increases was also different in Paired rats compared to Unpaired rats, as indicated by a significant Session × Pairing interaction (F(7.086,255.088) = 5.946, p < 0.001, ε = 0.709). Bonferroni-corrected post-hocs indicated that Paired rats made more US port entries than Unpaired rats for sessions 1-7 (p ≤ 0.027). However, for sessions 8-11, there was no significant difference (p = 0.126-0.701). This may be because Paired rats became more efficient in responding, making only one US port entry per trial or remaining in the port after the CS, while Unpaired rats may have learned to detect US deliveries by the sound of the syringe pump or by monitoring the port more closely. Again, there was no significant effect of Colony (F(1,36) = 0.003, p = 0.959) or Pairing × Colony interaction (F(1,36) = 0.491, p = 0488). There was also no significant Session × Colony interaction (F(7.086,255.088) = 1.125, p = 0.347, ε = 0.709) or Session × Pairing × Colony interaction (F(7.086,255.088) = 1.112, p = 0.356, ε = 0.709). Thus, while Paired rats learned to retrieve the sucrose US sooner than Unpaired rats did, by the end of autoshaping, both groups learned to retrieve the sucrose US, and there were no effects of breeding colony on this response.

ITI port entries varied slightly across the course of autoshaping (Figure 1f). Mixed-design ANOVA indicated that there was a significant overall decrease across Sessions (F(5.448,196.14) = 8.363, p < 0.001, ε = 0.545). However, ITI port entries did not differ according to Pairing (F(1,36) = 1.144, p = 0.292) or Colony (F(1,36) = 0.341, p = 0.563) and no Pairing × Colony interaction (F(1,36) = 1.56, p = 0.22). There was also no significant Session × Pairing interaction (F(5.448,196.14) = 1.64, p = 0.145, ε = 0.545), Session × Colony interaction (F(5.448,196.14) = 1.963, p = .08, ε = 0.545), or Session × Pairing × Colony interaction (F(5.448,196.14) = 0.847, p = 0.526, ε = 0.545). Thus, an overall decrease in ITI port entries suggests some habituation to the behavioural apparatus/procedure over the course of training, and there were no differences between breeding colonies or Paired and Unpaired rats in this response.

### Confirmatory analyses showed no effect of breeding colony on PCA score

PCA scores across autoshaping indicated that across experimental conditions, most animals were intermediates, with some tendencies towards goal-tracking, as evidenced by more negative PCA scores (Figure 2a). For Paired rats, a X^2^ test indicated there was no significant difference between breeding colonies in their final distribution across sign-tracking (K72 n = 1, R06 n = 1), goal-tracking (K72 n = 7, R06 n = 11), and intermediate phenotypes (K72 n = 7, R06 n = 3;X^2^(2) = 2.489, p = 0.288).

**Figure 2.**
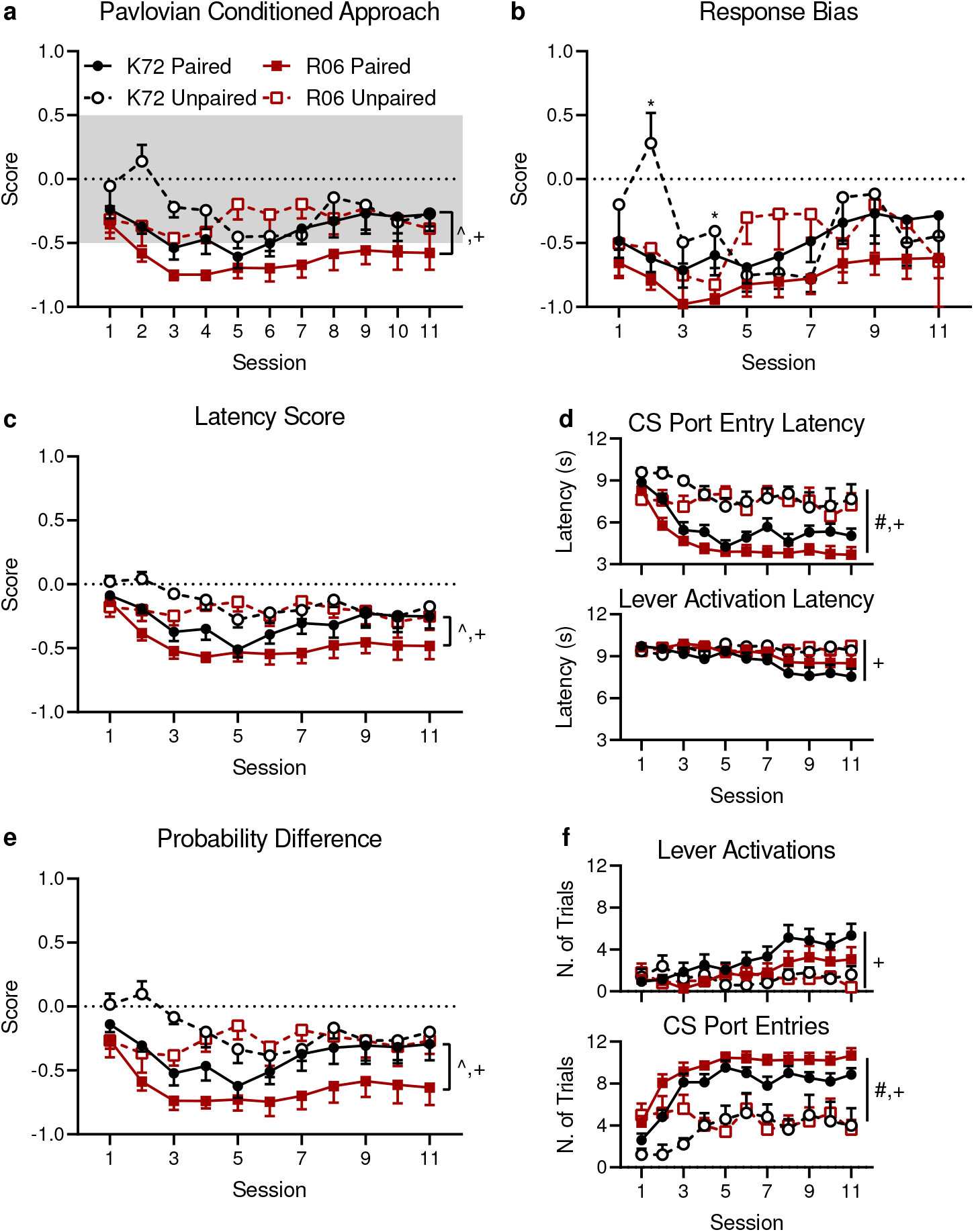
(a) PCA score indicated that our rats were mostly intermediates and goal-trackers. Confirmatory analyses demonstrated that Paired rats had more negative PCA scores than Unpaired rats did, indicating that Paired rats were more likely to approach the port while Unpaired rats did not show a clear preference for sign or goal-tracking. Linear mixed modelling of Paired rats alone also indicated that R06 rats goal-tracked more strongly than K72 rats did. Sign-trackers were defined by PCA scores ≥ 0.5 and PCA scores ≤ 0.5 defined goal-tracking. Intermediate PCA scores are shaded grey. (b) There were transient group differences in response bias (a measure of responding on the lever CS over the sucrose port), where on sessions 2 and 4, K72 rats showed more response bias compared to R06 animals, regardless of pairing condition (* p < 0.05 for Bonferroni-correct post-hoc comparisons). (c) There were no significant effects of breeding colony on latency score. (d) When analysing latency score components, R06 rats made significantly faster CS port entry latencies compared to K72 rats (top panel), but there was no effect of colony on lever activation latency (bottom panel). e) Similarly, there were no significant effects of breeding colony on probability difference. (f) When analysing probability difference components, there was no significant effect of colony on the number of trials with lever activations (top panel), but R06 rats made CS port entries on a greater number of trials than K72 rats did (bottom panel). + p < 0.05 for main effect of Pairing. K72 Paired n = 15, R06 Paired n = 15, K72 Unpaired n = 5, R06 Unpaired n = 5. # Main effect of colony on CS-triggered port entry latency and trials with CS-triggered port entries. ^ Main effects of colony on PCA score, latency score, and probability difference were observed when analysing Paired rats alone.

Confirmatory analyses of PCA score and its components did not support our hypothesis of significant differences between breeding colonies. While a mixed-design ANOVA suggested PCA score changed significantly across autoshaping Sessions (F(3.421,123.167) = 2.956, p = 0.029, ε = 0.342), there was only a significant difference between Paired and Unpaired rats (Pairing: F(1,36) = 4.963, p = 0.032). There was no significant main effect of breeding Colony (F(1,36) = 2.23, p = 0.144) or Pairing × Colony interaction (F(1,36) = 0.588, p = 0.448). There was also no significant Session × Pairing interaction (F(3.421,123.167) = 0.815, p = 0.502, ε = 0.342), Session × Colony interaction (F(3.421,123.167) = 1.513, p = 0.21, ε = 0.342), or Session × Pairing × Colony interaction (F(3.421,123.167) = 1.439, p = 0.231, ε = 0.342). Therefore, while this analysis demonstrated Paired rats had more negative PCA scores than Unpaired rats did, there was no effect of breeding colony on PCA scores across training.

Similar results were obtained when examining PCA score components. Response bias (Figure 2b), which measures how total responses are proportionally biased towards the lever CS (Figure 1a) over the sucrose port (Figure 1c), changed significantly across Sessions (F(4.329,155.852) = 3.566, p = 0.007, ε = 0.433). Paired and Unpaired rats did not significantly differ as there was no significant main effect of Pairing (F(1,36) = 1.856, p = 0.181). Overall, there was no significant difference between K72 and R06 rats (Colony: F(1,36) = 1.454, p = 0.236) and there was no significant Pairing × Colony interaction (F(1,36) = 0.411, p = 0.525). While there was a significant Session × Colony interaction (F(4.329,155.852) = 2.403, p = 0.047, ε = 0.433), Bonferroni-corrected post-hoc comparisons indicated that these differences were transient. K72 rats only had a significantly greater preference for the lever than R06 rats on sessions 2 (p = 0.004) and 4 (p = 0.031). Finally, there was no significant Session × Pairing interaction (F(4.329,155.852) = 1.318, p = 0.264, ε = 0.433) or Session × Pairing × Colony interaction (F(4.329,155.852) = 2.254, p = 0.061, ε = 0.433). Thus, across groups, pairing condition had no effect on response bias, and breeding colony had only a transient effect.

The latency score is a ratio which compares how quickly animals contacted the lever CS relative to how quickly they enter the sucrose port. As shown in Figure 2c, it followed the same trajectory as the overall PCA score. Latency score significantly varied across autoshaping, as shown by a significant effect of Session (F(2.793,100.537) = 3.7, p = 0.016, ε = 0.279). Paired rats approached the port more quickly than Unpaired rats did, as indicated by a significant Pairing effect (F(1,36) = 8.776, p = 0.005). However, there was no significant Session × Pairing interaction (F(2.793,100.537) = 1.373, p = 0.256, ε = 0.279). Overall latency scores between K72 and R06 rats were not significantly different (Colony: F(1,36) = 2.74, p = 0.107) and there was no Pairing × Colony interaction (F(1,36) = 0.635, p = 0.431). There was also no significant Session × Colony interaction (F(2.793,100.537) = 0.749, p = 0.517, ε = 0.279) or Session × Pairing × Colony interaction (F(2.793,100.537) = 0.707, p = 0.541, ε = 0.279). Latency score therefore followed the same pattern as PCA scores and showed that while Paired rats approached the sucrose port more quickly than Unpaired rats did, there were no significant effects of breeding colony.

The components of the latency score, latency to enter the port during CS presentation and latency to activate the lever CS, are shown in Figure 2d. Latency to, respectively, enter the port during CS presentation and activate the lever CS significantly varied across Sessions (Figure 2d, top panel; F(3.073,110.634) = 10.205, p < 0.001, ε = 0.307 and Figure 2d, bottom panel; F(3.124,112.463) = 2.966, p <= 0.033, ε = 0.312) and these changes were specific to Paired rats (Figure 2d, top panel; Session × Pairing: F(3.063, 110.634) = 3.238 = 0.024, ε = 0.307 and Figure 2d, bottom panel; F(3.124,112.463) = 3.012, p = 0.031, ε = 0.312). In response to CS presentation, Paired rats approached the port significantly faster than Unpaired rats did (Pairing: F(1,36) = 41.925, p < 0.001), as did R06 rats compared to K72 rats (Colony: F(1,36) = 4.504, p = 0.041). However, neither Pairing nor Colony influenced the latency to activate the lever CS (Pairing: F(1,36) = 2.174, p = 0.149; Colony: F(1,36) = 0.448, p = 0.507). There were no other significant effects on the latency to enter the port CS (Session × Colony: F(3.073,110.634) = 0.994, p = 0.4, ε = 0.307; Pairing × Colony: F(1,36) = 0.496, p = 0.486; Session × Pairing × Colony: F(3.073,110.634) = 0.958, p = 0.417, ε = 0.307) or latency to activate the lever CS (Session × Colony: F(3.124,112.463) = 0.65, p = 0.59, ε = 0.312; Pairing × Colony: F(1,36) = 0.409, p = 0.526; Session × Pairing × Colony: F(3.124,112.463) = 0.312, p = 0.824, ε = 0.312). Therefore, while latency score overall did not significantly vary with breeding colony, compared to K72 rats, R06 rats entered the port faster after CS presentation.

The third component of the PCA score, probability difference, measures how much more likely animals are to interact with the lever CS on any given trial than enter the sucrose port. Probability difference (Figure 2e) significantly varied across Sessions (F(3.16,113.749) = 2.965, p = 0.033, ε = 0.316), following the same pattern as PCA score and its other components. Similarly, Paired rats were less likely to make lever activations than Unpaired rats were (Pairing: F(1,36) = 7.748, p = 0.009), but there were no significant overall differences between K72 and R06 rats (Colony: F(1,36) = 2.863, p = 0.099). There was also no significant Pairing × Colony interaction (F(1,36) = 0.772, p = 0.385), Session × Pairing interaction (F(3.16,113.749) = 1.053, p = 0.374, ε = 0.316), Session × Colony interaction (F(3.16,113.749) = 1.022, p = 0.388, ε = 0.316), or Session × Pairing × Colony interaction (F(3.16,113.749) = 1.064, p = 0.369, ε = 0.316). Thus, probability difference also followed the same pattern as PCA scores and showed that Paired rats were more likely than Unpaired rats to enter the sucrose port but there were no effects of breeding colony.

The number of trials with lever activations and trials with CS-triggered port entries, which comprise the probability difference, are shown in Figure 2f. Trials with lever activations and CS-triggered port entries varied across Session (Figure 2f, top panel; F(3.394,122.173) = 2.957, p = 0.029, ε = 0.339 and Figure 2f, bottom panel F(3.83,137.863) = 8.845, p < 0.001, ε = 0.383, respectively) and these changes were specific to Paired rats (Session × Pairing: Figure 2f, top panel F(3.394,122.173) = 3.332, p = 0.017, ε = 0.339 and Figure 2f, bottom panel F(3.83,137.863) = 3.761, p = 0.007, ε = 0.383). Compared to Unpaired rats, Paired rats had significantly more trials with CS-triggered port entries (Figure 2f, top panel; F(1,36) = 58.283, p < 0.001) but did not have more trials with lever activations (Figure 2f, bottom panel F(1,36) = 1.907, p = 0.176). There was no significant effect of breeding colony on lever activations (Figure 2f, top panel; F(1,36) = 0.652, p = 0.425), but R06 rats had significantly more trials with CS-triggered port entries than K72 rats did (Figure 2f, bottom panel; F(1,36) = 4.871, p = 0.034). There were no other significant effects for the number of trials with lever activations (Figure 2f, top panel; Session × Colony: F(3.394,122.173) = 0.835, p = 0.489, ε = 0.339; Pairing × Colony: F(1,36) = 0.465, p = 0.5; Session × Pairing × Colony: F(3.394,122.173) = 0.563, p = 0.662, ε = 0.339) or the number of trials with CS port entries (Figure 2f, bottom panel; Session × Colony: F(3.83,137.863) = 1.823, p = 0.131, ε = 0.383; Pairing × Colony: F(1,36) = 0.562, p = 0.458; Session × Pairing × Colony: F(3.83,137.863) = 1.321, p = 0.266, ε = 0.383). Consistent with the colony effects observed on latency scores (Figure 2d), these results showed that R06 rats were more likely to make CS-triggered port entries compared to K72 rats.

### Linear mixed modelling of Paired rats alone revealed colony effects

Rather than use an ANOVA, Fitzpatrick and colleagues reported using a linear mixed model to analyse differences Pavlovian conditioned approach behaviour between Charles River rats from different breeding colonies [9]. Using this approach, they found significant colony effects [9]. We therefore followed their approach and analysed Paired rats alone using linear mixed models. Only Paired rats showed evidence of acquiring Pavlovian conditioned approach behaviour, with low levels of responding in Unpaired rats, as expected. Therefore, excluding the Unpaired rats from statistical analyses removed animals that neither sign-nor goal-tracked and that are therefore strictly irrelevant to the question of whether breeding colony affects sign- and goal-tracking. We produced a series of models (Table 4) and found that the best model (Model 1), which was 3.16 times more likely than the next best model, indicated that breeding colony was a significant predictor of PCA score.

**Table 4.** Linear mixed modelling results

| Model | AICc | $\Delta i$ | wi | ER |
| --- | --- | --- | --- | --- |
| 1. Colony + Session | 4.172 | 0 | 0.759 | - |
| 2. Session | 6.473 | 2.301 | 0.240 | 3.16 |
| 3. Colony + Session +<br>Colony × Session | 17.176 | 13.004 | 0.001 | 666.47 |
| 4. Colony | 24.976 | 20.804 | $2.30 \times 10^{-5}$ | 32925.41 |
| 5. Null | 27.149 | 22.977 | $7.77 \times 10^{-6}$ | 97587.04 |

Model 1 indicated that there were effects of breeding colony, with overall PCA scores of K72 rats 0.219 ± 0.1 points higher than PCA scores of R06 rats. The effects observed in the present study were also comparable with those reported by Fitzpatrick and colleagues [9]. Calculating η*p*^2^ from our tests of fixed effects (F(1,38.265) = 4.778, p = 0.035) produced a 90% CI = [0.004, 0.271], which overlapped with the results reported by Fitzpatrick and colleagues (F(1,132.21) = 10.69, p = 0.001, 90% CI = [0.018, 0.154]) [9]. In the present study, the effect of Session (F(10,261.972) = 4.49, p < 0.001, 90% CI = [0.059, 0.182]) was also comparable to Colony. There was also a high correlation between PCA scores across ses-sions(AR1 ρ = 0.82 ± 0.028, Wald Z = 28.93, p < 0.001). These results indicate that breeding colony had an effect on PCA score that was similar in magnitude to that of training sessions and comparable to the results reported by Fitzpatrick and colleagues [9].

The difference between the results of our confirmatory analyses and linear mixed modelling were due to the inclusion of Unpaired rats masking the effects of breeding colony in our confirmatory analyses. Linear mixed modelling also reveals how comparable the effects in the present study are to prior work by Fitzpatrick and colleagues [9]. However, the present results are not due to the use of linear mixed modelling per se. Similar effects can be observed when using mixed-design ANOVA to analyse Pavlovian autoshaping data in Paired rats alone. When using this statistical approach, we observe significant main effects of Colony on ΔCS port entries (F(1,28) = 4.716, p = 0.039), unadjusted CS port entries (F(1,28) = 6.976, p = 0.013), probability difference (F(1,28) = 5.559, p = 0.026), latency score (F(1,28) = 5.023, p = 0.033), and PCA score (F(1,28) = 4.476, p = 0.043). Therefore, it appears that the effect of breeding colony was masked by the inclusion of the Unpaired rats in confirmatory analyses, because when Paired rats are analysed alone K72 and R06 rats differ on multiple measures.

### No effect of breeding colony on conditioned reinforcement

After autoshaping, the conditioning chambers were reconfigured so that the sucrose port was replaced with the lever CS and flanked on either side with two new nosepoke ports. We then assessed the extent to which the lever CS had acquired conditioned reinforcing properties by determining whether the rats would nosepoke to earn presentations of the CS alone, with no sucrose US. As shown in Figure 3a, there was not clear evidence for discrimination between active and inactive nosepokes (F(1,36) = 3.804, p = 0.059). This suggests that the cue had no value as a conditioned reinforcer. Overall, Paired rats made more nosepokes than Unpaired rats did (Pairing: F(1,36) = 5.3, p = 0.027; Figure 3a), but there was no indication that this was specific to the active nosepoke (Nosepoke × Pairing: F(1,36) = 1.84, p = 0.183). Similarly, there was no significant effect of breeding Colony (F(1,36) = 2.268, p = 0.141), Pairing × Colony interaction (F(1,36) = 0.415, p = 0.523), Nosepoke × Colony interaction (F(1,36) = 0.496, p = 0.486), or Nosepoke × Pairing × Colony interaction (F(1,36) = 0.126, p = 0.725). Thus, these findings suggest that across pairing condition and breeding colonies, rats did not attribute conditioned reinforcing value to the lever CS.

**Figure 3.**
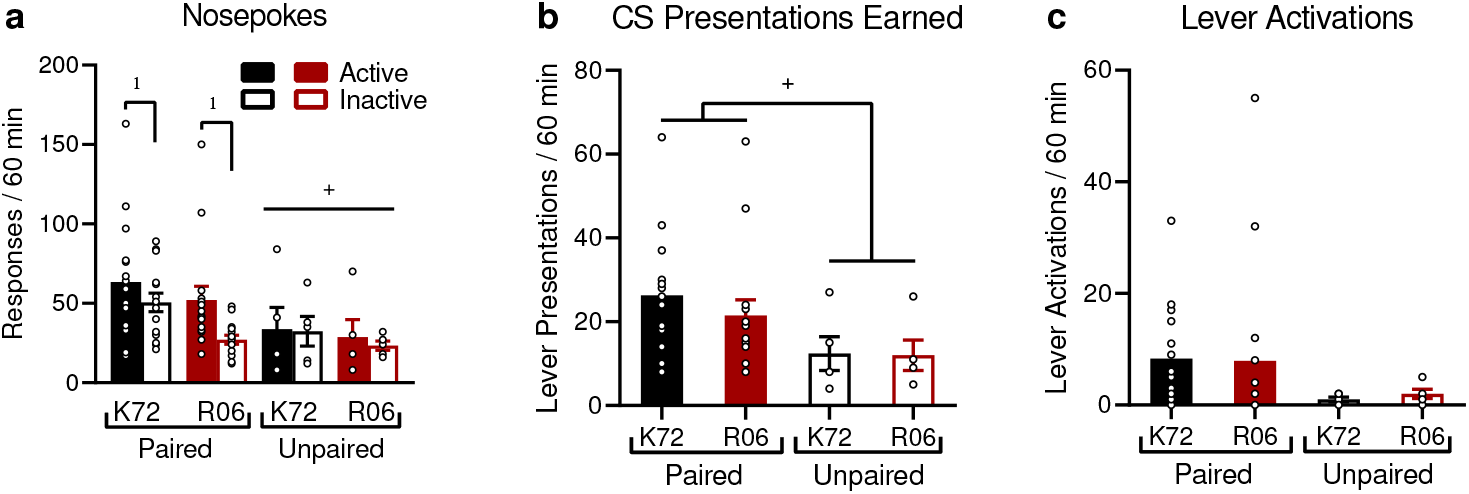
Conditioning chambers were reconfigured so that rats could nosepoke to earn presentations of the lever CS alone, without the sucrose US. (a) Paired rats nosepoked more than Unpaired rats did. When analysed alone, Paired rats showed nosepoke discrimination, responding more on the active vs. inactive nosepoke (as indicated by *). This indicates that in these rats, the lever CS acquired conditioned reinforcing properties. However, there were no specific effects of breeding colony. (b) Paired rats earned more CS presentations than Unpaired rats did (as indicated by +), but there were no effects of breeding colony. (c) While Paired rats appear to interact more with the lever, this was not statistically significant. K72 Paired n = 15, R06 Paired n = 15, K72 Unpaired n = 5, R06 Unpaired n = 5.

Because the lack of significant discrimination between the active vs. inactive ports was surprising, we conducted an exploratory analysis of Paired rats alone to investigate whether they showed significant operant responding for conditioned reinforcement. A mixed-design ANOVA with Paired rats alone showed that these animals did favour the active nosepoke (F(1,28) = 10.113, p = 0.004; Figure 3a). While K72 rats made marginally more nosepokes than R06 rats did (F(1,28) = 4.21, p = 0.0497), this was not selective for the active nosepoke (Nosepoke × Colony interaction: F(1,28) = 1.038, p = 0.317). These results suggest that while the lever CS did have value as a conditioned reinforcer for Paired rats, there was no effect of breeding colony.

Confirmatory analyses showed no effect of breeding colony on the number of CS presentations earned (Figure 3b). A 2×2 ANOVA revealed that while Paired rats overall earned more presentations of the lever CS compared to Unpaired rats (Pairing: F(1,36) = 5.692, p = 0.022), there was no significant effect of Colony (F(1,36) = 0.283, p = 0.598) or Pairing × Colony interaction (F(1,36) = 0.202, p = 0.655). This indicates that Paired rats earned more CS presentations than Unpaired rats did, but there was no effect of breeding colony.

Once the lever CS was presented, there were no significant differences in the rate at which rats from different groups activated it (Figure 3c). While inspection of Figure 3c suggests that Paired rats might interact more with the lever CS, this was not statistically significant (Pairing: F(1,36) = 2.606, p = 0.115). There was also no significant effect of Colony (F(1,36) = 0.005, p = 0.942) or Pairing × Colony interaction (F(1,36) = 0.029, p = 0.866). Thus there were no effects of breeding colony on lever activations during the conditioned reinforcement test.

## Discussion

The present study empirically examined whether Long-Evans rats from different breeding colonies operated by the same vendor differed in Pavlovian conditioned approach behaviour. Our confirmatory analyses found little evidence of differences in PCA phenotypes, but effects may have been masked by the inclusion of the Unpaired group, for which CS presentation was not paired with sucrose delivery. When Paired rats were analysed alone, using the same linear mixed modelling approach used in prior studies [9], the findings demonstrate an effect of breeding colony that was comparable to previous studies that have found colony differences [9]. In parallel, we found that breeding colony had no significant effects on operant responding for a CS, a test for the CS’s conditioned reinforcing properties.

### When the lever CS was paired with sucrose, rats acquired Pavlovian conditioned approach behaviour

Over the course of 11 autoshaping sessions, we compared responding to the CS in Paired versus Unpaired rats and found that Paired rats showed significant conditioned responding to the CS. Paired rats significantly increased their rate of ΔCS port entries across sessions, while Unpaired rats made close to no ΔCS port entries after 11 sessions. Similarly, Paired—but not Unpaired—rats increased their rate of goal-tracking responses across sessions. This included increases in both the latency to enter the port after the onset of CS presentation, and in the number of trials with a CS-triggered port entry. Our evidence for the acquisition of sign-tracking is more equivocal, with only 2 rats classified as sign-trackers (and a third with PCA score = 0.499). There was only a marginal effect of pairing on the increase in lever CS activations across sessions (Figure 1a, Session × Pairing interaction effect). However, across sessions, Paired rats activated the lever CS faster than Unpaired rats did (Figure 2d), and Paired rats also had a greater number of CS trials with lever activations (Figure 2f). Thus, Paired rats acquired Pavlovian conditioned approach responses, with most animals tending towards goal-tracking, but with evidence for sign-tracking behaviour as well.

### Paired rats from K72 sign-tracked more

Our preregistered confirmatory analyses did not provide support for our hypotheses that K72 rats would have higher PCA scores or be more likely to be classified as sign-trackers. However, the inclusion of Unpaired rats could have masked differences between K72 and R06 rats. This is due to both the lower-than-expected level of sign-tracking in the present cohort and the calculation of PCA scores as an average of ratios (Table 3). This floor effect in sign-tracking made the Unpaired rats appear more similar to Paired rats in PCA score and component measures and added variability that may have masked the effects of breeding colony in the subsequent analyses. Unpaired rats were included as a control to demonstrate that CS-triggered conditioned approach responding requires that the CS and US be paired in time. As such, low levels of responding to the CS were expected in Unpaired rats [24, 25]. This low level of responding in Unpaired rats was consistent across the entire experiment, with very few lever activations, port entries, or nosepokes throughout the 11 autoshaping sessions and the conditioned reinforcement test session. Excluding the Unpaired rats from the analyses therefore removed the added and uninformative variability of animals for which we have no theoretical or empirical reason to expect sign- or goal-tracking behaviour from. Moreover, the calculation of PCA scores and its components as ratios can be easily skewed by low rates of responding. This is because an Unpaired rat that makes two or three responses can have a similar PCA score to a rat that makes a dozen responses, yet we would not claim that the Unpaired rat in this scenario has acquired sign- or goal-tracking to the same extent as the Paired rat has. Therefore, when we analysed the Paired rats alone, following the approach used by Fitzpatrick and colleagues [9], we found results similar to theirs. Specifically, K72 rats had significantly higher PCA scores than R06 rats did, supporting our hypothesis that K72 rats would sign-track more.

Moreover, this effect of breeding colony is not simply due to using a statistical approach different from our planned ANOVA analyses. If we use mixed-design ANOVAs to analyse Pavlovian autoshaping data in the Paired rats alone, we also observe significant main effects of Colony on CS-triggered conditioned approach behaviour, as measured by ΔCS port entries, unadjusted CS port entries, probability difference, latency score, and PCA score. Therefore, although our confirmatory analyses did not support our hypotheses that K72 rats would have higher PCA scores, our exploratory analyses indicate that these effects are significant but were masked by the inclusion of the Unpaired groups. Although these analyses do not indicate, in the present study, that breeding colony produces significantly different final phenotype frequencies, these analyses did experimentally replicate the large analyses performed by Fitzpatrick and colleagues across several research projects and experimenters [9].

### No breeding colony effects on conditioned reinforcement

In addition to assessing potential colony differences in Pavlovian conditioned approach behaviour, we also examined differences in the instrumental pursuit of the CS. Determining whether rats will spontaneously acquire and perform a new instrumental response to obtain presentations of a CS alone, without its associated US, is a critical test for conditioned reinforcement [60–62]. Thus, after PCA training, the rats were allowed to nosepoke into a new port for presentations of the CS alone. If rats nosepoke significantly more into the active versus inactive port, this would confirm that the CS has acquired conditioned reinforcing properties. The inclusion of Unpaired rats in analysis of the data from the conditioned reinforcement test also masked nosepoke discrimination. When we analysed nosepokes in accordance with our preregistered analysis plan (that is, including both Paired and Unpaired rats), we found no significant discrimination between nosepoking into the active vs. inactive port. This was expected for Unpaired rats, for whom the lever CS should have no conditioned reinforcing properties, because it was never explicitly paired with sucrose. When we re-analysed these data with Paired rats only, we found a significant preference for the active port, demonstrating that the lever CS was a conditioned reinforcer in Paired rats. Therefore, while Paired rats did attribute conditioned reinforcing properties to the lever CS, our hypothesis that K72 rats would show greater instrumental pursuit of a CS than R06 rats would was not supported.

### Vendor differences and contributing factors

The present study makes an important contribution to the literature on vendor differences because it examines the less well-characterized effects of different breeding colonies from the same vendor. Fitzpatrick and colleagues did demonstrate within-vendor effects by analysing across many different research projects and experimenters [9]. This approach has merit, because it involves a very large sample size. However, there are also drawbacks due to potential confounding factors. For example, between-experimenter effects have been reported in rodents, such as olfactory analgesia associated with human male experimenters [63]. There are also some reports of seasonal or circannual effects in rats and mice, but these effects are not always large or consistent [64, 65]. The present study controlled for these potential confounds by direct empirical testing of within-vendor breeding colony effects, holding experimenter and time of year constant across experimental groups. To our knowledge (Table 2), this is one of only two papers to empirically test whether there are differences in behavioural phenotype between rats sourced from different breeding colonies operated by the same vendor.

It is likely that both genetic and environmental factors contribute towards potential vendor breeding colony differences in Pavlovian conditioned approach behaviour. Fitzpatrick and colleagues demonstrated that there were genetic differences in rats from different breeding colonies that were associated with their autoshaping phenotypes [9]. A study by Gileta and colleagues has recently underscored the importance of genetic factors in PCA and showed significant differences in Sprague Dawley rats within and between vendors, including between the Raleigh (R06) and Kingston (K72) Charles River facilities [66] we compared here. Environmental differences may involve subtle physical or operational differences between facilities that influence animal behaviour. Environmental factors may also involve transport between breeding facilities and the research institute, which has been shown is stressful for animals [67, 68]. Moreover, it is assumed that transporting animals a longer distance will result in greater levels of stress [69], which would suggest that depending on geographical location, rats from one breeding facility can be subjected to greater transport stress than rats from another. This is a plausible explanation since stressed animals have been shown to interact less with a Pavlovian cue [70]. In support, we observed that animals from the more distant colony (R06) appeared to goal-track more than rats from the less distant facility (K72). Within-vendor breeding colony differences may therefore be due to a combination of genetic and environmental factors. Single-housing could also be a factor in this effect, as single housing of animals can be associated with poor welfare [71]. Single-housing was used here to remain consistent with previous studies [12], including our unpublished studies.

### Limitations

The present study specifically examined breeding colony differences in Pavlovian conditioned approach behaviour in male rats and cannot be used to identify the factors that might contribute to such differences. It is also notable that the effects size of colony was lower than expected, which resulted in the masking of colony effects by the inclusion of the Unpaired rats. The exclusion of the Unpaired rats from our analyses also suggests that caution is required in interpreting the results of these analyses. This being said, pairing vs. unpairing lever CS and sucrose presentation significantly influenced responding. In Paired rats that had received lever CS presentations explicitly paired with sucrose, the CS came to evoke conditioned approach responses. For example, compared to Unparied rats, Paired rats made more CS-triggered port entries, had lower latencies to CS-triggered port entries and lever activations, and were more likely to make CS-triggered port entries and lever activations. Future studies might use a double-lever design, where an inert CS-lever is tested alongside a US-paired, CS+ lever in the same rats, foregoing the need for an Unpaired control group [12, 70]. A double-lever approach would not follow the classic single-lever designs used in key studies [8, 9] but would overcome some of the statistical limitations from using separate Unpaired control animals. Finally, male rats have predominantly been used in Pavlovian conditioned approach studies [8, 9, 12] and the majority of the literature on vendor differences also uses male rats (Tables 1 and 2). As such, we used male rats for comparison with this literature. In this context, it is important to replicate these findings in female rats.

The dominance of the goal-tracking and intermediate phenotypes in the present study also suggest some limitations with respect to drawing conclusions about sign-tracking. While PCA scores exist on a continuum and K72 rats sign-tracked more or had more positive PCA scores than did R06 rats, we cannot draw conclusions about sign-tracking per se. A reasonable alternative interpretation of our findings is therefore that they say less about the overall balance between sign- and goal-tracking and more about a reduction in goal-tracking in a group of animals that are biased towards goal-tracking. In this case, further studies with cohorts that are more balanced or more predisposed towards sign-tracking are required.

### Conclusions

Following observations of potential within-vendor differences across several Pavlovian conditioned approach experiments, we performed a direct empirical study to test the hypothesis that breeding colony influences autoshaping phenotype (i.e., sign-vs. goal-tracking in response to CS presentation). While our confirmatory analyses did not support our hypothesis, this appeared to be due to the inclusion of Unpaired rats in our design, which masked the effects of breeding colony in our statistical analyses. Indeed, analysing Paired rats alone replicated previous findings that breeding colony influences PCA score. However, we also found that breeding colony did not significantly alter the proportion of sign- or goal-trackers nor did it produce differences in the conditioned reinforcement value of the lever CS. We therefore suggest that experimenters studying Pavlovian autoshaping should be alert to the possibility of breeding colony effects and take care to record the specific facilities supplying animals for each experiment.

## Disclosures

### Author Contributions

Conceptualization – SYK, AU, NC; Data curation – SYK; Formal analysis – SYK, AU; Funding acquisition – SYK, AU, NC, ANS; Investigation – SYK, AU; Methodology – SYK, AU, NC; Project administration – SYK, NC; Resources – NC; Software – SYK; Supervision – SYK, NC, ANS; Validation – SYK; Visualization – SYK; Writing – original draft – SYK, ANS; Writing – review & editing – SYK, ANS.

## Acknowledgements

The authors dedicate this paper to the memory of Dr Nadia Chaudhri.

